# RPPA survey of cancer hotspot panel proteins and cell markers in matched tumor-normal human breast and kidney samples reveals a weak correlation between proteins expression and public transcriptome repositories

**DOI:** 10.1101/2023.11.09.566387

**Authors:** Krisztina Paal, Csaba Konrad, Noemi Karnok, David Bui, Christos Chinopoulos

**Affiliations:** Department of Biochemistry, Semmelweis University, Budapest 1094, Hungary; Feil Family Brain and Mind Research Institute, Weill Cornell Medicine, New York, NY 10065, USA

**Keywords:** Reverse phase protein array, oncogene, tumor-suppressor gene, hotspot, proteomics

## Abstract

Despite the tremendous global effort in discovering and cataloguing somatic mutations in cancer-related genes and the impact they exert on human cancers, the effects of these mutations on the levels of protein expression are inadequately addressed. Here, we semi-quantitated the expression of 48 (out of 50) proteins represented by the most widely used cancer hotspot panel, the Ion AmpliSeq™ Cancer Hotspot Panel v2, and 14 cell markers commonly used as ‘house-keeping’ proteins, using reverse phase protein array (RPPA) technology in 586 human breast- and 192 kidney matched tumor-normal samples. We further correlated our RPPA data with gene chip data from the Gene Expression Omnibus repository fetched through the TNMplot portal. Nearly half of proteins expression exhibited a negative correlation from their gene expression, while several of the remaining were only moderately correlated. Our data complement the vast information obtained elsewhere by transcriptomic analysis of the respective genes and gene products, potentially assisting in harmonizing proteomic with NGS outputs.

## INTRODUCTION

All cancers harbor highly heterogeneous somatic mutations ^1^ that accumulate during progression of the disease ^2^. Most mutations are random and inconsequential -known as passenger mutations ^3, 4^-while others contribute to tumor initiation, progression and maintenance, referred to as driver mutations ^5, 6^. Driver mutations may bear greater impact if present in oncogenes and/or tumor suppressor genes ^7^. More recently, the number of mutations per megabase (mut/Mb) irrespective of their status as driver or passenger -referred to as tumor mutational burden (TMB)-has emerged as a biomarker, specifically in terms of sensitivity of the harboring tumor to immune checkpoint inhibitors^8^. ‘Hotspot’ mutations are those that occur in tumor samples significantly more frequently than expected from a background frequency characterized by genes, cancer types, mutation types and sequence contexts ^9^. The explosion in computing power, tissue banking resources and collaborations among institutions across the globe observed in the past few decades have led to the assembly of compendia analyzing whole genomes of thousands of primary tumors and hundreds of metastases or local recurrences ^6^ as well as to the development of numerous computational methods for detecting and cataloguing mutations and even predicting hotspots ^10^. Such efforts have culminated in the emergence of mega-catalogues of somatic mutations in cancer, such as COSMIC (Catalogue Of Somatic Mutations In Cancer) ^11^. COSMIC covers all the genetic mechanisms by which somatic mutations promote cancer, including non-coding mutations, gene fusions, copy-number variants and drug-resistance mutations; furthermore, one subproject -COSMIC-3D-offers predictions on variables regarding the expressed protein, such as protein structure (folding, rigidity), charge, stability, ligand binding properties or confidence in maintaining protein-protein interaction interfaces, all with implications on draggability ^12^. Having said that, it is important to emphasize that any information offered regarding effect of a mutation on protein level by COSMIC-3D is only *a prediction*, and is not an experimentally-verified result. Finally, in the Gene Expression Omnibus (GEO) database managed by the National Center for Biotechnology Information (NCBI) ^13^, high-throughput screening genomics data derived from microarray or RNA-Seq experiments can be found. In there, comparisons can be made concerning the levels of mRNA expression between healthy-and tumorous tissues for a variety of cancers. The user-friendly, freely available TNMplot web tool (accessible at TNMplot.com) can be used for a visual representation of GEO queries and highly-filtered, structured data downloads regarding expression levels of selected genes can be obtained.

On the other hand, in 2013, the Cancer Proteome Atlas (TCPA) emerged from MD Anderson ^14^, a publicly available collection of cancer functional proteomics data with parallel DNA and RNA data https://tcpaportal.org/tcpa/; its most updated version is 3.8 (March 2022) ^15, 14, 16^. Although this is an extremely useful and comprehensive platform integrating RPPA with TCGA output, it only offers data from tumor material and there are no samples from healthy tissues (nor matched, neither non-matched).

Comparisons in protein expression of cancer-related proteins between cancerous *vs* healthy tissues appears only sporadically in Formalin-Fixed Paraffin-Embedded (FFPE) samples where immunohistochemical assays were exclusively used; among the 48 cancer-related proteins that we investigated, we only found tumor-normal protein expression comparisons for FGFR3 ^17^, SMARCB1 ^18^ and TP53 ^19, 20^; quantitation of any IHC results harbors a low degree of confidence. Data on additional proteins can be found in the human protein atlas https://www.proteinatlas.org/ but they also suffer from the same limitation, *i.e.* they are obtained from IHC and thus non-readily quantifiable. Here, by using reverse phase protein array (RPPA) technology we semi-quantitated the expression of 48 out of 50 proteins represented by the most widely used cancer hotspot panel, the Ion AmpliSeq™ Cancer Hotspot Panel v2, in 586 matched tumor-normal human breast and 192 kidney samples; furthermore, we assessed the expression of 14 cell markers commonly used as ‘house-keeping’ proteins ^21^. We finally correlated our RPPA data with gene chip data from the Gene Expression Omnibus repository filtered for paired tumor and adjacent normal tissues, obtained through the TNMplot portal. Our data show a weak correlation between protein-and gene expression, not only for the those represented by the cancer hotspot panel, but also the cell markers. Finally, our efforts establish the workflow for the RPPA analysis of tumor-suppressor and oncogene proteins otherwise mostly sought for mutations in their encoding genes in a medium-range pool size of matched, tumor-normal samples.

## MATERIALS AND METHODS

Samples: 586 human breast- and 192 kidney matched tumor-normal samples were provided by Voronezh State University, which were collected from nearby clinics in the Voronezh area in the time interval 2017-2020. The samples were provided under the auspice of the bilateral, transnational project 2017-2.3.4-TET-RU-2017-00003. All samples were frozen to -20 °C within 1 hour of excision and shipped in dry ice in bar-coded cryovials. Ethical permission for this project has been obtained and is on permanent display in https://tinyurl.com/rppahuethicalperm (file number: 35302-5/2017/EKU). Personal data of the patients from whom the samples originated were alphanumerically encrypted using a 128-bit encryption algorithm and stored in a secure server.

RPPA: The workflow for RPPA was the following (detailed protocols can be found in http://rppa.hu/Protocols.html):

i. solubilization of samples to lysates and protein amount determination: Samples (20-200 mg) were suspended in a buffer (named “Benzonase buffer”), the composition of which was: 20 mM Trizma, 2 mM MgCl_2_, protease inhibitors (EDTA-free, Thermo #A32955), Benzonase (Pierce #88700), 0.2 mM BeSO_4_ and 5 mM NaF, pH 7.2 (HCl). After a few minutes while samples were kept on ice, they were homogenized by bead beating in a Precellys 24 homogenizer (Bertin Technologies SAS) using zirconium oxide 2 mm beads. Subsequently, the homogenized samples were incubated at 37 °C while shaking @500 rpm. After 5 min, homogenates were spun @ 3,500 rpm for 2.5 min. Subsequently, an equal amount of volume of a 2x Laemmli buffer (named “Laemmli buffer”) was added, the composition of which was: 4% SDS, 20% glycerol, 120 mM Trims, 4 mM dithiothreitol (DTT), 5 mM EDTA, 5 mM EGTA, pH 6.8 (HCl) and the bead-beating protocol was repeated once. Then, tubes were incubated at 97.5 °C for 10 min, while shaking @500 rpm. Lysates were subsequently transferred to new tubes avoiding the top-lipid fraction and any remaining insoluble material, spun at 12,700 rpm @RT for 10 min. Supernatants were collected and diluted 10-fold using a 1-to-1 mixture of Benzonase buffer and Laemmli buffer. Subsequently, protein concentration was determined by IR spectrometry measuring amide bond absorbance using a Direct Detect IR Spectrometer (Merck KGaA). If protein concentration (of the undiluted lysate) was > 21 mg/ml, a calculated amount of buffer was added (to decrease concentration to < 21 mg/ml) and lysate protein concentration was re-estimated. All lysates were set to 1-20 mg/ml, aliquoted in new bar-coded cryovials and stored @ -80 °C until use.
ii. Lysates loading to 96-well plates and then to 384-well plates: Lysates were thawn while rocking and then loaded first in 96-well plates and then to 384-well plates using an automated epMotion 5073 liquid handling system (Eppendorf, AG). Dilutions were performed iteratively so that each lysate is greater or equal than 2.05 mg/ml. At the end of each robotic run, each lysate was present in 1x, 2x, 4x, 8x and 16x fold dilutions in 4 replicates each, per 384-well plate. Well plates were then bar-coded, sealed and kept @ -80 °C until use.
iii. Spotting of lysates using an Arrayer: Spotting of the lysates occurred by transferring 1 nL of lysate from the 384-well plate (all dilutions and replicates) on Oncyte SuperNOVA nitrocellulose bar-coded slides (Grace Bio-labs) twice per slide using an Aushon 2470 arrayer with 16 pins (Aushon, Billerica, MA, USA). 13-15 well plates and 50 slides were loaded in the arrayer per run, thus, each slide was spotted by 9,660 spots. Each spotting run lasted ∼29 hours. To counter batch effects, samples were randomly assigned into two, partially overlapping runs. Because the high number of samples the distribution of protein loaded should be the same between the runs. We observed significant batch effects for some antibodies, therefore the runs were normalized to have equal mean values for each antibody between the two runs; supplementary figures 1C (unnormalized) and 1D (normalized) show histograms of RPPA read intensities for Vimentin, which was an example of an antibody with a strong batch effect. Slides were kept in thermo-controlled incubators @15 °C until use.
iv. Staining of slides: Slides were manually stained in incubation chambers (Grace Bio-labs) using a quadruple amplification protocol that allowed using extremely low titers of antibodies, see tables 2 and 3. Specifically: slides were re-hydrated with Tris-buffered saline containing 0.1 % Tween-20 for 10 minutes. Subsequently, they were ‘blocked’ with 5% bovine serum albumin (Sigma, A7906) and 0.1 % Tween-20 for 1 hour. Slides were subsequently treated with Avidin (immediately after the block step without washing) followed by Biotin, followed by 0.3% H_2_O_2_ and several wash steps in-between and before applying the primary (tables 2 and 3) and secondary antibodies (these were either donkey anti-mouse or donkey anti-rabbit, depending on the host organism where the primary antibody was raised; titers of secondary antibodies were 1:10,000 for both). All antibodies were pre-tested by western blotting using the same quadruple amplification protocol described above or pre-validated by the RPPA platform of Institute Curie (IC, https://institut-curie.org/) and see tables 2 and 3. Afterwards, slides were treated with an amplification module (Bio-Rad, cat # 1708230) followed by applying streptavidin conjugated with IRDye 800 (1:2,000) and several wash steps in-between. In addition, one slide per RPPA run was stained by FCF Fast Green (Sigma F7258) for normalization of spots during the imaging process (see below). Slides were scanned as described below within a few hours after the last washing step and following a short drying protocol by spinning at 300 rpm for 5 min.
v. Slide/blot imaging and spot quantification: Slides were loaded (24 slides at a time, using an autoloader) in an Innoscan 710-IR (Innopsys Inc). The FCF-stained slide was scanned at 670 nm, and the IRDye 800-stained slides at 785 nm, using the GAL file generated by the Arrayer. Spot quantification and output to excel was automatically performed using MAPIX software (Innopsys).

Antibody validation: Antibodies were selected according to the following criteria: (i) reactivity to human tissues, (ii) monoclonality (raised in rabbits or mice) or recombinant, (iii) monospecificity (in Western blot), and if applicable, (iv) the epitope should not be located in hotspot area. On the basis of the above, > 400 commercially available antibodies were pre-screened and ∼100 selected and tested. Antibody monospecificity was verified using standard Western blotting, using 7 random breast- and 7 random kidney samples. Blots were imaged using an Azure 600 Imaging System. 48 antibodies were deemed as suitable for RPPA. Some were pre-validated by the RPPA platform of Institute Curie (https://institut-curie.org/), see also tables 2 and 3. For PIK3CA and SMO proteins, no suitable antibodies were found. Detailed information regarding all antibodies can be found in db.rppa.hu portal (registration required; contact the corresponding author C.C.).

Data analysis: Normalization of RPPA data was done by the protein stain (FCF) data using a custom-made python script (https://github.com/csabak/RPPA/tree/main). First, background was subtracted and then the exponential function y = A * exp(K * x) + C was fitted to the protein standard of each sample, where y is the fluorescence value (Fprot), x is the dilution ratio and A, K and C are fitted parameters (see also supplementary figure 1B). Then, background was subtracted from the antibody values and all values below 3000 intensity were marked as low reads and discarded. Each antibody read was normalized by the calculated FCF value by the calibration curve for each dilution, and Fprot values with z-scores above 3 and below -3 were discarded.

Plots of RPPA and gene chip data (the latter obtained from the TNMplot portal https://tnmplot.com using paired tumor and adjacent normal tissues as described in ^22^) were generated by SuperPlotsOfData web app ^23^; volcano plots were generated by the VolcaNoseR web app ^24^. Slope-colored paired RPPA data shown in the supplementary figures 2-7 were generated in ggplot 2 using a custom-made R script deposited at https://github.com/JoachimGoedhart/Slopes-colored.

## RESULTS

Semi-quantification of proteins expression represented by the Ion AmpliSeq™ Cancer Hotspot Panel v2 and cell markers in matched tumor-normal human breast and kidney samples by RPPA

All samples were provided by Voronezh State University, which were collected from nearby clinics in the Voronezh area in the time interval 2017-2020. Breast tumor and adjacent normal tissue were obtained from 293 patients aged 29-83 years; kidney tumor and adjacent normal tissue were obtained from 96 patients aged 33-84 years. Their age distribution is shown in the violin plots of supplementary figure 1A. Each sample (after being converted to lysate, as described under the section of RPPA workflow in the Materials and Methods) was probed 40 times using 1x, 2x, 4x, 8x, and 16x dilution ratios in 4×2 replicates, as shown in supplementary figure 1B. In supplementary figure 1B, the term “0R7Z4MA_002_T_1” is an alphanumerically encrypted tumor sample. From each such curve, an fprot value was obtained from each sample, and plotted quasi-randomly as shown in the left panels (using fprot in the y-axes in figures 1-6 (figures 1, 2 and 3 for breast samples, oncoproteins, tumor suppressor proteins and cell markers respectively, and figures 4, 5 and 6 for kidney samples oncoproteins, tumor suppressor proteins and cell markers respectively). In each panel, the mean and +/-1*SD is included (black lines). Data parity (healthy-tumor samples from the same patient connected by a line) is shown in the supplementary figures 2-7. Figure panels of the supplementary figures 2-7 for breast and kidneys are superimposed to the proteins shown in main figures 1-6 for rapid visual queuing. In the main figures data some extreme outliers have been arbitrarily removed (>5 SD) so as to scale down the figure for better visual comparisons; however all data points are present in the corresponding data pair (matching tumor-normal) in the supplementary material figures. We have selected to probe for proteins corresponding to the Ion AmpliSeq™ Cancer Hotspot Panel v2 because this is the most commonly employed panel among datasets, grants, patents, clinical trials, and policy documents, see table 1. With this item 2,856 mutations can be probed (https://tinyurl.com/IonAmpliSeqmutations), distributed among 50 (proto)oncogenes and tumor suppressor genes. We have been able to find RPPA-suitable antibodies for 48 out of these 50 proteins [for proteins APC, p16INK4A, VEGFR2, NPM1, RET, SMARCB1 and SRC two suitable antibodies were found, designated as (AB1) and (AB2)]. No suitable antibody was found for PIK3CA and SMO proteins. Name of proteins, their aliases, abbreviation, short description, Research Resource Identifiers (RRIDs) of antibodies used, epitopes and titers are shown in table 2.

**Legend to figure 1:**
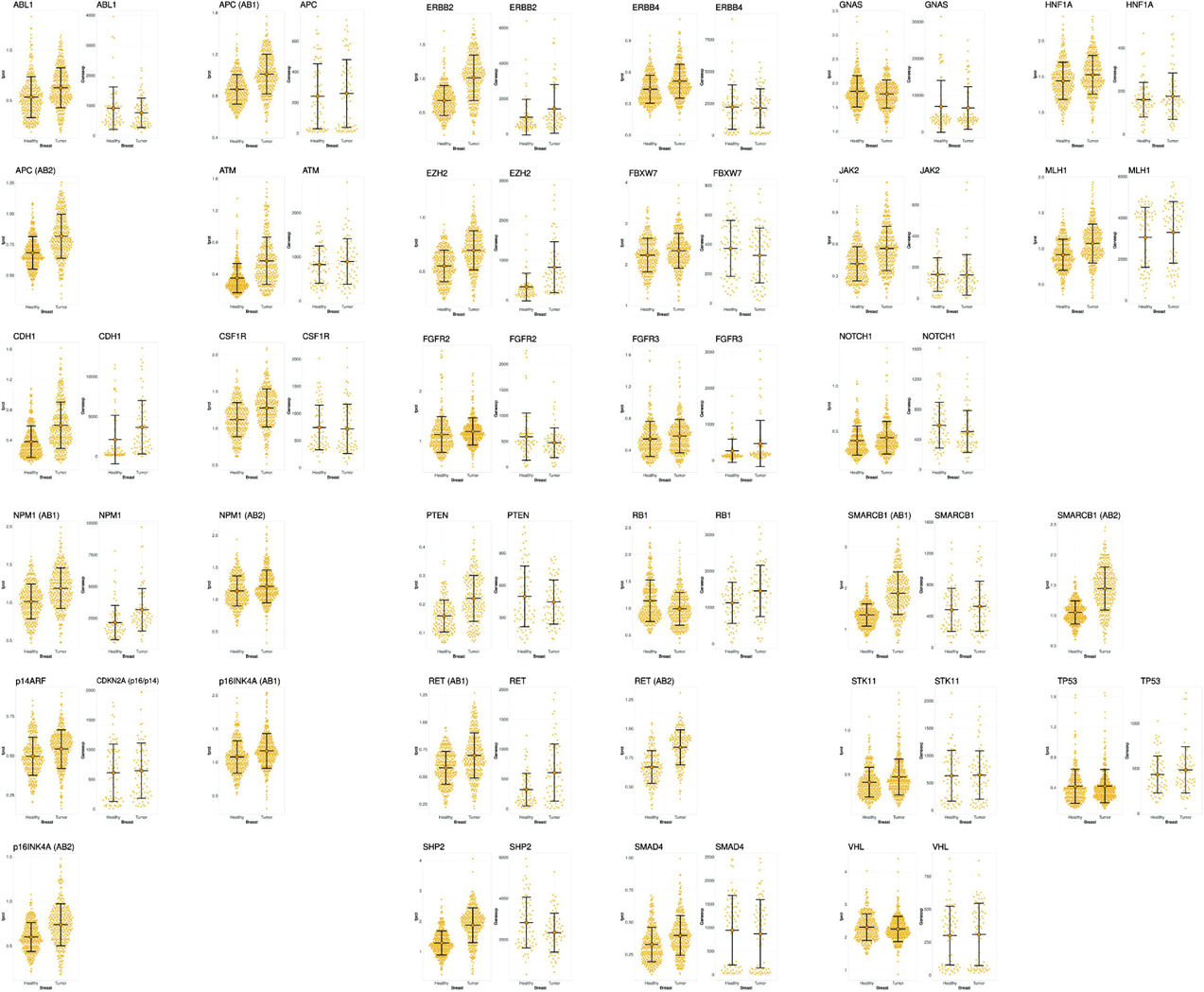
Quasi-random distribution of expression levels of oncoproteins (fprot, left panels) and respective oncogenes (Geneexp, right panels) in human breast matched tumor-normal samples. Black lines indicate mean +/-1 SD. (AB1) and (AB2) signify the type of the antibody used for the respective protein.

**Legend to figure 2:**
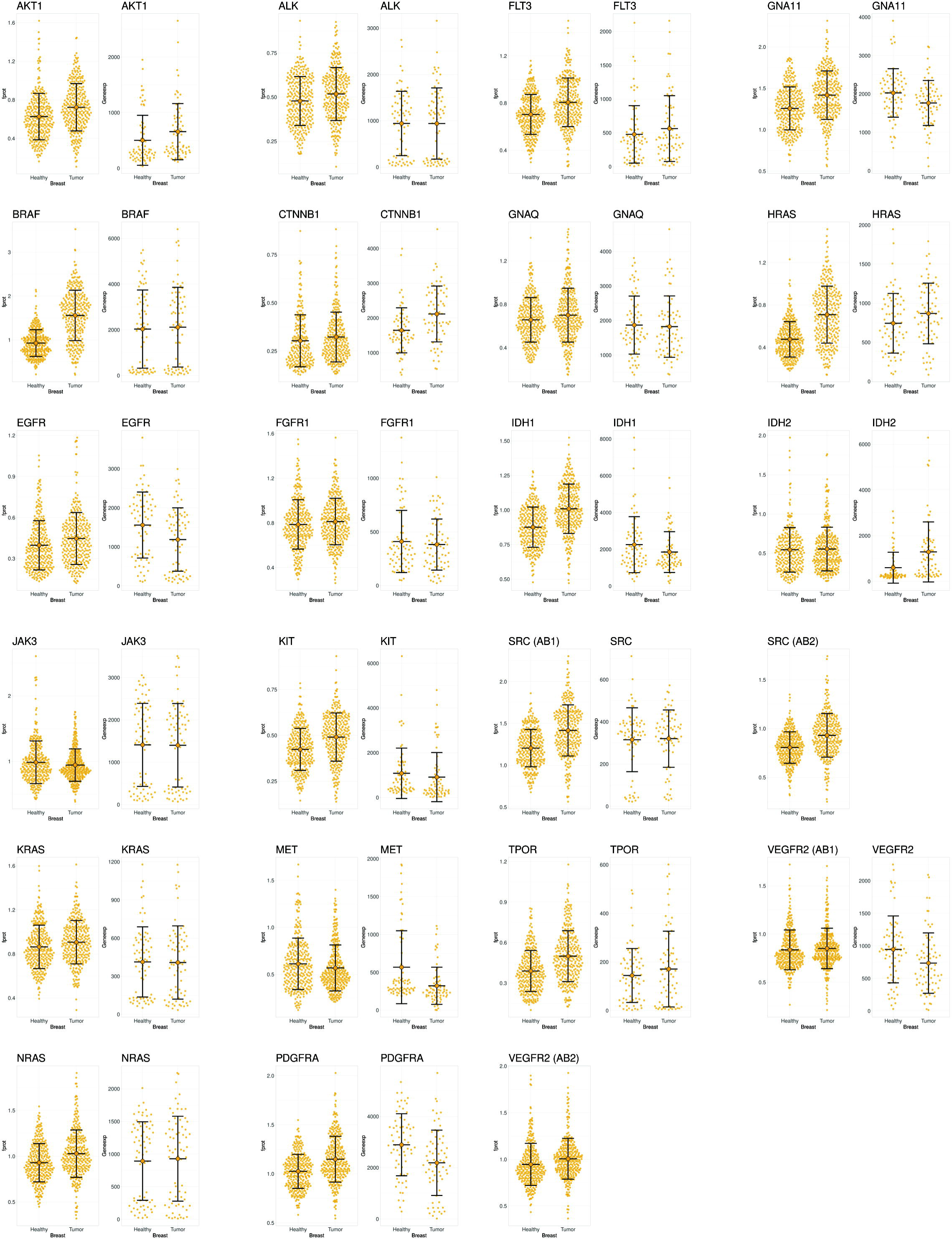
Quasi-random distribution of expression levels of tumor suppressor proteins (fprot, left panels) and respective tumor suppressor genes (Geneexp, right panels) in human breast matched tumor-normal samples. Black lines indicate mean +/-1 SD. (AB1) and (AB2) signify the type of the antibody used for the respective protein.

**Legend to figure 3:**
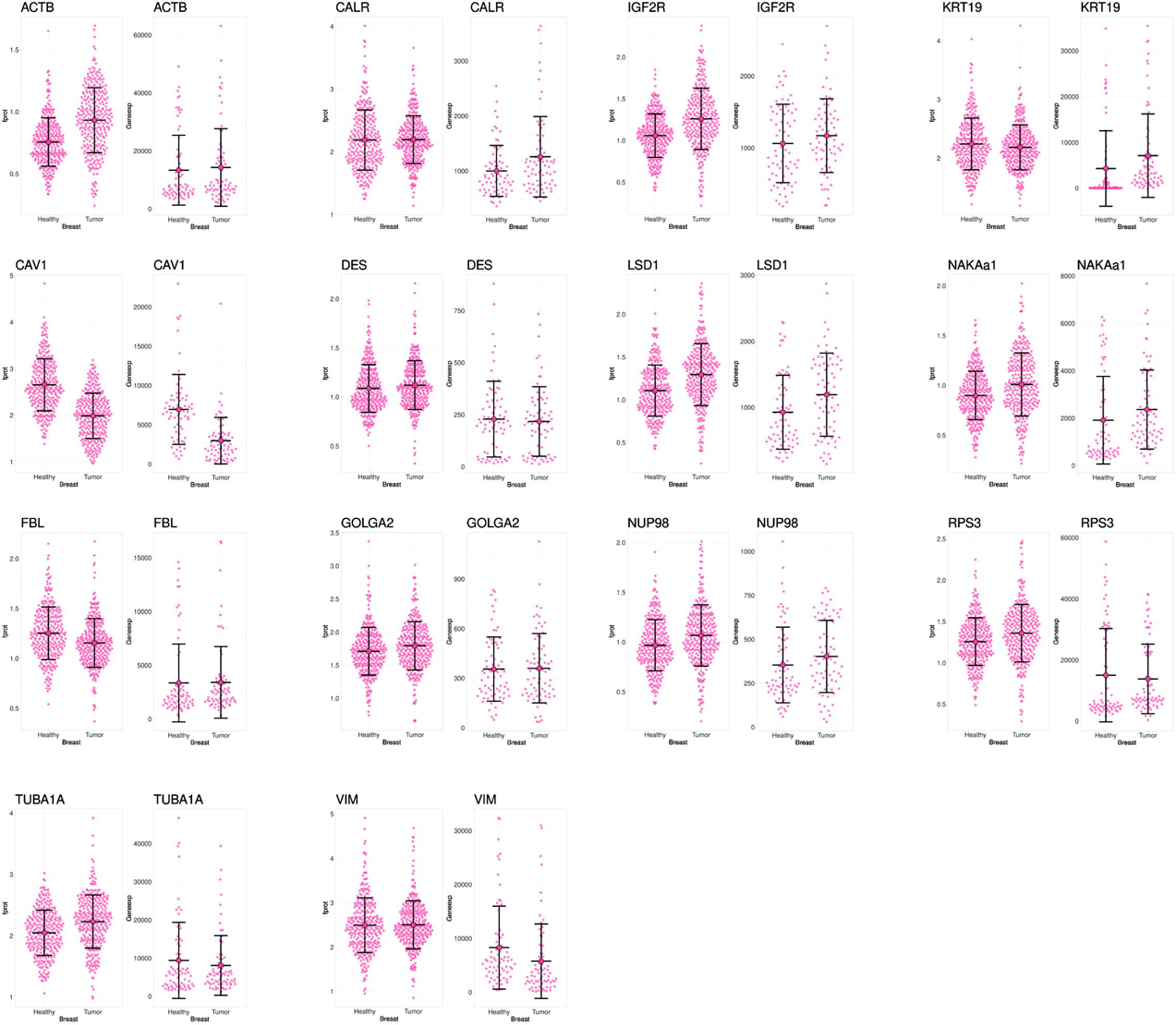
Quasi-random distribution of expression levels of cell marker proteins (fprot, left panels) and respective cell marker genes (Geneexp, right panels) in human breast matched tumor-normal samples. Black lines indicate mean +/-1 SD.

**Legend to figure 4:**
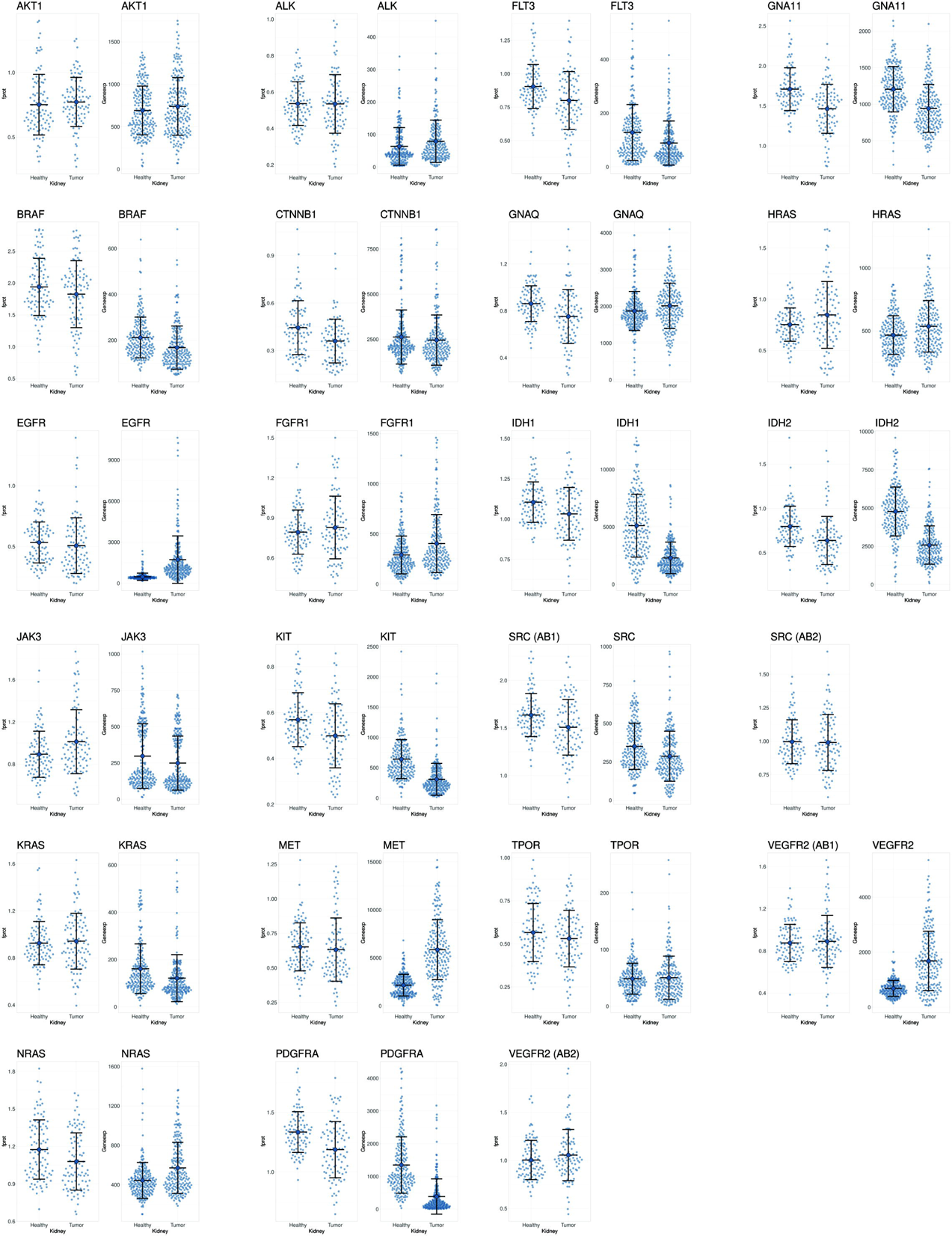
Quasi-random distribution of expression levels of oncoproteins (fprot, left panels) and respective oncogenes (Geneexp, right panels) in human kidney matched tumor-normal samples. Black lines indicate mean +/-1 SD. (AB1) and (AB2) signify the type of the antibody used for the respective protein.

**Legend to figure 5:**
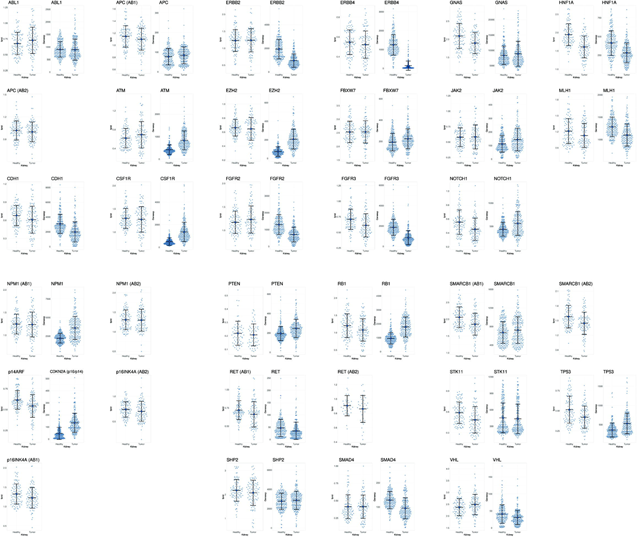
Quasi-random distribution of expression levels of tumor suppressor proteins (fprot, left panels) and respective tumor suppressor genes (Geneexp, right panels) in human kidney matched tumor-normal samples. Black lines indicate mean +/-1 SD. (AB1) and (AB2) signify the type of the antibody used for the respective protein.

**Legend to figure 6:**
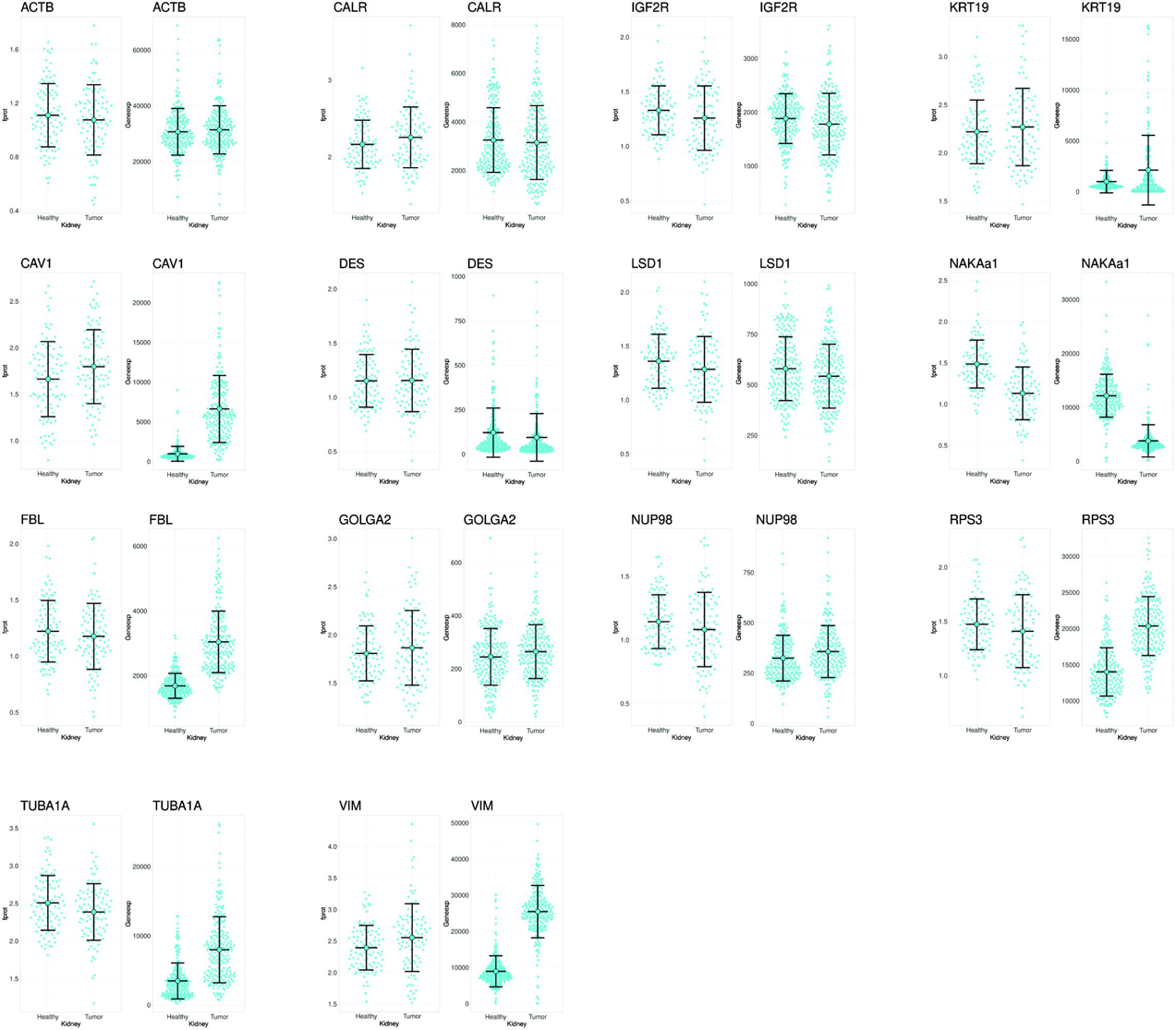
Quasi-random distribution of expression levels of cell marker proteins (fprot, left panels) and respective cell marker genes (Geneexp, right panels) in human kidney matched tumor-normal samples. Black lines indicate mean +/-1 SD.

Arbitrarily, we have divided these 48 proteins in oncoproteins and tumor-suppressor proteins to minimize figure panels clutter, even though several of them exhibit both functions depending on conditions ^25, 26, 27^. We labeled the following ones as oncoproteins: AKT1, ALK, BRAF, CTNNB1, EGFR, FGFR1, FLT3, GNA11, GNAQ, HRAS, IDH1, IDH2, JAK3, KIT, KRAS, MET, NRAS, PDGFRA,SRC, TPOR, VEGFR2. Those labelled as tumor suppressor proteins were: ABL1, APC, ATM, CDH1, CSF1R, ERBB2, ERBB4, EZH2, p16INK4A, p14ARF, FBXW7, FGFR2, FGFR3, GNAS, HNF1A, JAK2, MLH1, NOTCH1, NPM1, PTEN, RB1, RET, SHP2, SMAD4, SMARCB1, STK11, TP53, VHL. Finally, we used antibodies for these house-keeping proteins: ACTB, CALR,CAV1, DES, FBL, GOLGA2, IGF2R, KRT19, LSD1, NAKAa1, NUP98, RPS3, TUBA1A, VIM. Name of cell markers, abbreviations, RRIDs of antibodies used and their titers is shown in table 3. The cell markers were selected to represent a most wide repertoire of cellular functions/compartments as described previously ^21^.

Statistical significance of proteins expression as a function of fold change is depicted in the volcano plots in figure 7: proteins expressed in breast samples are addressed in the top row volcano plot (proteins coded by oncogenes, tumor suppressors or cell markers, as indicated in the panels) while for kidney samples in the bottom row of volcano plots (same arrangements, as indicated in the panels). As shown in figure 7, only MET protein exhibited a statistically significant decrease in expression in breast samples (from the oncoproteins + tumor suppressor proteins). From the cell markers, only FBL, GOLGA2, KRT19, DES, VIM and CALR would be suitable for normalization. CAV1 exhibited an extremely significant decrease in expression in tumor vs healthy breast tissues. On the other hand, no oncoproteins nor tumor suppressor protein exhibited a statistically significant decrease in the kidney tumor vs healthy tissues; concerning the cell markers, the α subunit of the Na+/K+ ATPase (NAKAa1) showed an extremely significant decrease in expression in tumor vs healthy kidney tissues. All other markers exhibited no statistically significant differences. It is also notable that there is good agreement among almost all of the proteins against which two antibodies were used exemplified by close proximity (except NPM1), affording additional confidence as to the RPPA workflow (compare SMARC (AB1) with SMARC (AB2), APC (AB1) with APC (AB2), p16INKA (AB1) with p16INKA (AB2) and RET (AB1) with RET (AB2); other proteins against which two antibodies were used are not labeled in the volcano plots because they did not reach statistical significance (grey points).

**Legend to figure 7:**
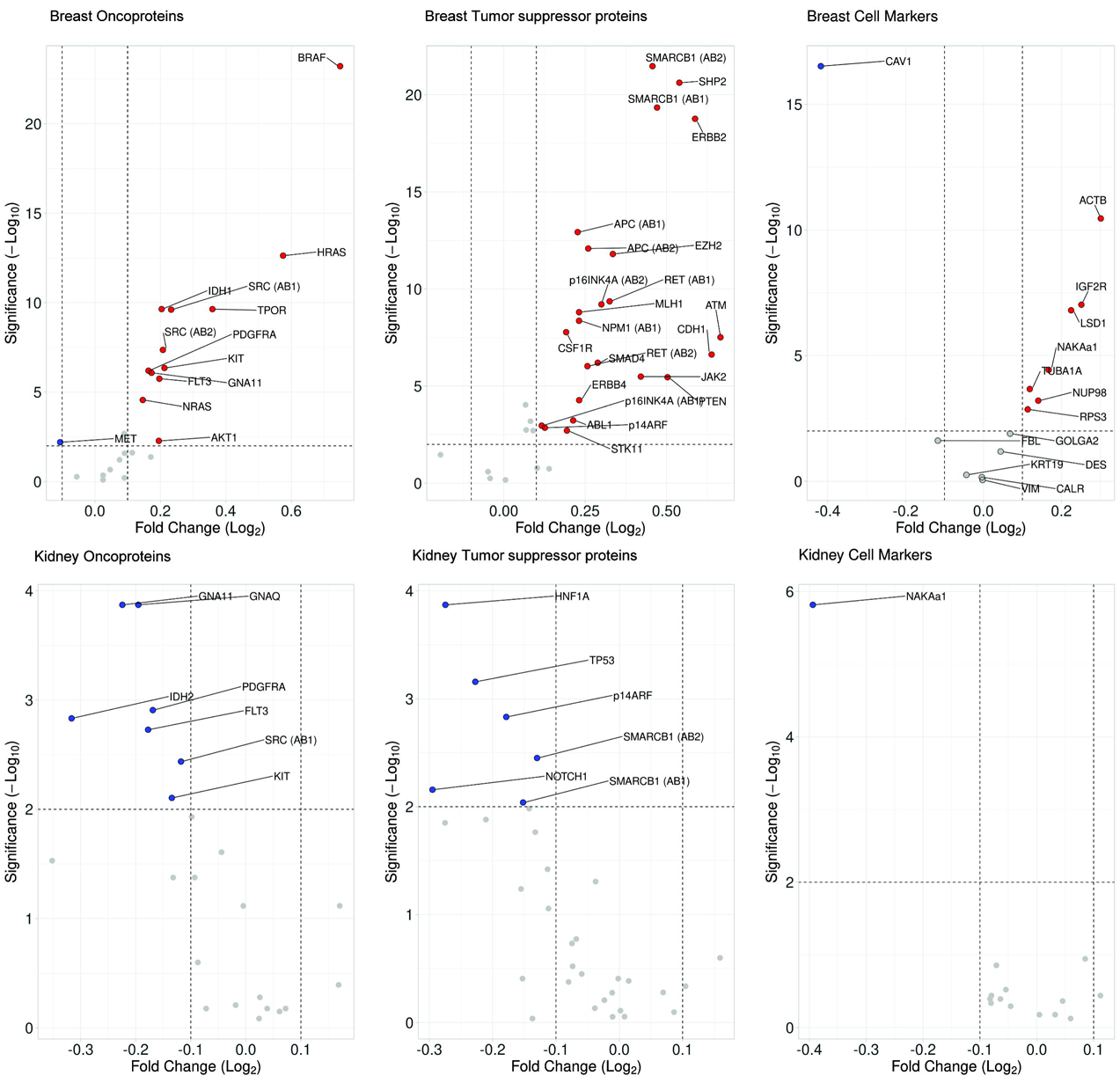
Volcano plot of breast (top row) and kidney (bottom row) oncoproteins, tumor suppressor proteins and cell markers depicting statistical significance versus magnitude of change. Fold changes and paired t-tests were run on the complete breast and kidney datasets, followed by correction for multiple comparisons using the Benjamini-Hochberg method to adjust for false discovery rate. Cutoffs were set to p-adjusted < 0.05 and log_2_ fold change > ± 0.1. The data was divided into oncoproteins, tumor suppressors, and cell markers for clearer visualization.

Comparison of the expression of genes represented by the Ion AmpliSeq™ Cancer Hotspot Panel v2 and cell markers in matched tumor-normal human breast and kidney samples using fetched data from NCBI-GEO Gene chip data fetched from NCBI-GEO through the TNMplot portal are depicted in the right panels of the main figures 1-6 (y-axes labeled as “Geneexp”). In each panel, the mean and +/-1*SD is also shown in black lines. Whenever two antibodies were suitable (against APC, p16INK4A, VEGFR2, NPM1, RET, SMARCB1 and SRC), the corresponding gene chip data is shown only to the right of RPPA data using AB1. It must be emphasized that these data are from paired healthy-tumor data repositories, but are not related to the samples that we used for RPPA.

### Weak correlation between RPPA- and gene chip data

As shown in figures 1-6, there is a discord between gene and protein expression for the same gene, for several gene and gene products. Due to the lack of gene expression data of our samples and the varying sample distributions, we only compared the numerical value of the increase or decrease in tumorous samples (relative to the healthy samples) between the transcriptional and the translational level, in order to illustrate degree of similarity. We made the calculation for both the median and the mean as neither of them were singly suitable to represent the sample population in all cases, see figure 8. However, as such, it must be emphasized that the numerical comparison is disproportionately sensitive in cases where very small changes are compared. Cases where the gene and protein expression change reliably in the same direction to the same degree were scarce. However, there are many gene-protein pairs where both measures shifted in the same direction, though to different extent, as shown in figure 8. In other cases, we observed a complete disassociation in detected transcriptional and translational changes, discussed below.

**Legend to figure 8:**
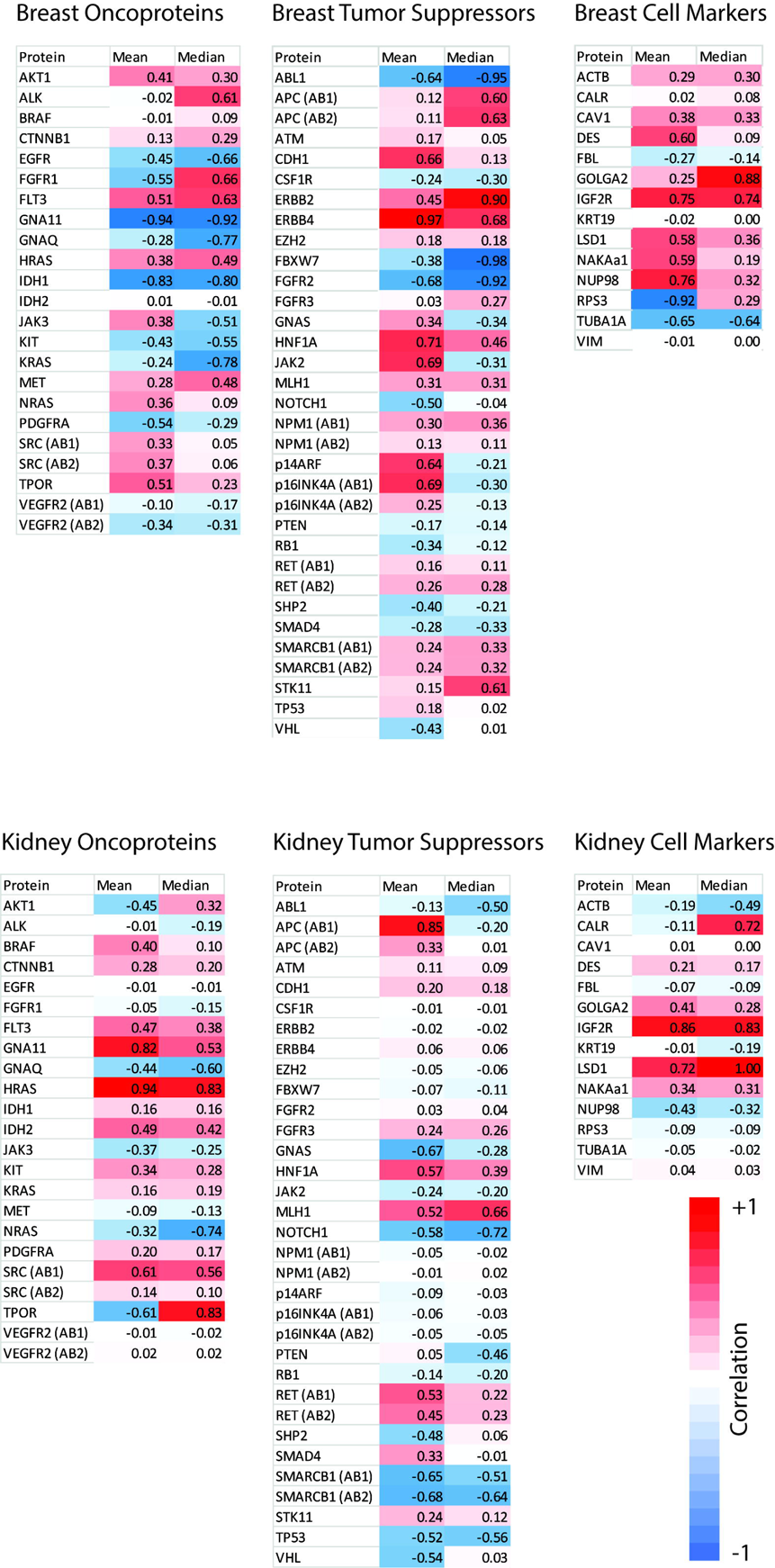
The similarity between the numerical values of the mean or median relative tumorous protein amount and mean or median relative tumorous gene expression. The scale can be described as 1 for equal, 0 for unrelated, -1 for inverse.

### Principal Component Analysis and Uniform Manifold Approximation and Projection of RPPA data

Further RPPA data visualization was done by 2D projecting by means of Principal Component Analysis (PCA) and Uniform Manifold Approximation and Projection (UMAP) using their python implementations (scikit-learn and umap respectively) for breast samples. The number of the kidney samples were not sufficient for such kind of analysis. For the breast datasets, each protein was normalized to have zero mean and equal variance of 1. Algorithms were run on four subsets: 1) using all the data, 2) excluding the cell markers, 3) oncoproteins only, 4) tumor suppressor proteins only. The four PCA runs had explained variances of 55-65% by the first two components, indicating that important details might be lost in this projection. A slight separation between tumor and healthy samples is seen on the breast data when using tumor suppressor proteins exclusively (figure 9A). Nonlinear structure was better preserved by UMAP. While no clear separation of tumor and healthy samples was observed, there is some clustering of breast samples using the three datasets where cell markers were excluded (figure 9B-D).

**Legend to figure 9:**
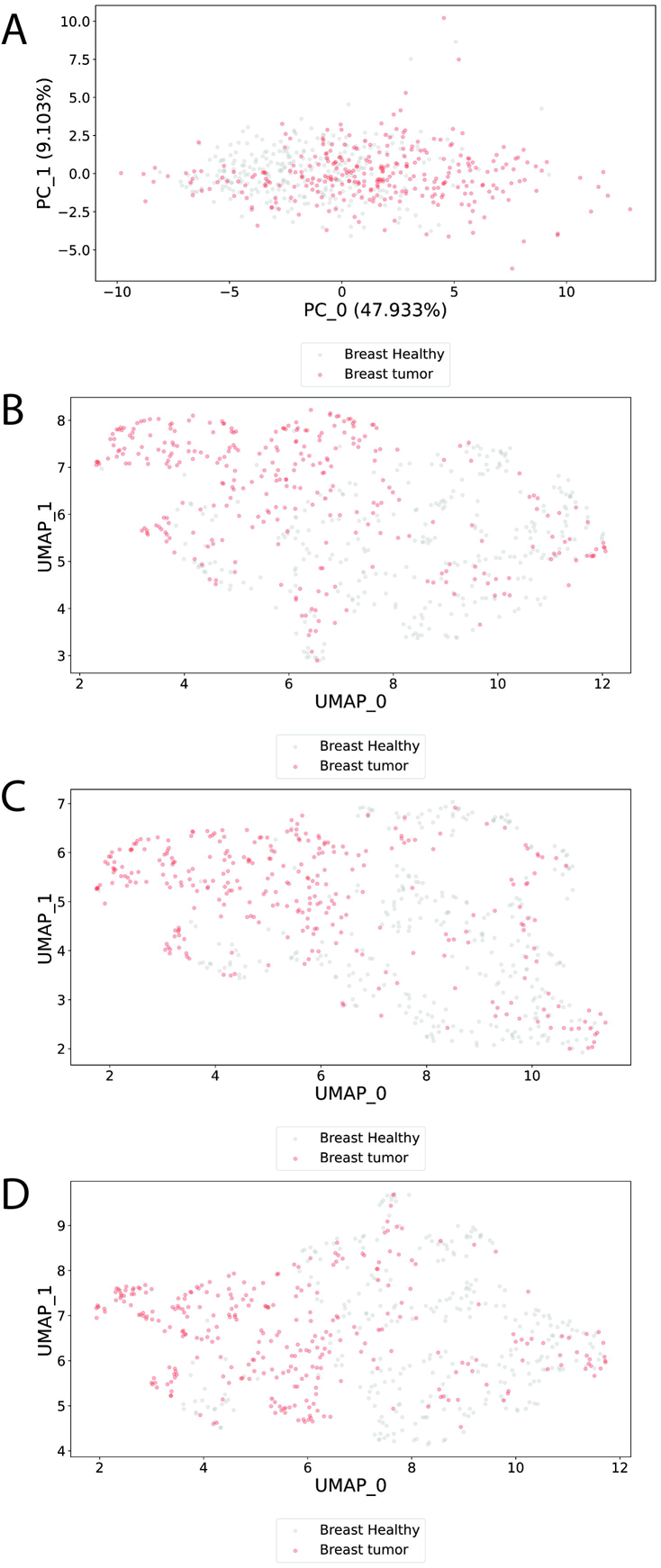
Principal Component Analysis (A) and Uniform Manifold Approximation and Projection (B-D) of RPPA data. Gray dots represent healthy-while red dots represent tumor samples. For PCA (panel A), retained variance along the two principal components is indicated on the axes. A includes data from tumor suppressor proteins in breast samples. B, C, and D include data from oncoproteins plus tumor suppressor proteins, oncoproteins alone, and tumor suppressor proteins alone, respectively.

## DISCUSSION

In the present work, 48 out of 50 of the proteins represented by the most widely used cancer hotspot panel, the Ion AmpliSeq™ Cancer Hotspot Panel v2 were semi-quantitated using reverse phase protein array (RPPA) technology in 586 human breast- and 192 kidney matched tumor-normal samples. We further probed for 14 cell markers commonly used as ‘house-keeping’ proteins, because of the controversial nature of total protein quantification by conventional means between healthy and tumorous tissues. Tumorous tissues exhibit very high lipid content, DNA and RNA (adding viscosity), necrotic segments and calcifications, rendering sample lysis and protein determination difficult. The method that we used here for protein determination prior to RPPA (IR spectroscopy of amide bonds) is devoid of some confounding factors most commonly afflicting conventional methods such as Biuret and bicinchoninic acid assay related to false positivity due to reactivity of non-protein components; however, protein extraction and determination of cancer tissues such as those from breast or kidney still carries a significant risk for under- and over-estimations. Thus, the normalization of samples (apart from the FCF stain of the spotted slides per RPPA run) to some other ‘house-keeping” protein is of critical importance. Yet, as we show here and in agreement with prior literature ^21^, only some of the cell marker proteins were similarly expressed in healthy-vs the tumor of the organ (FBL, GOLGA2, KRT19, DES, VIM and CALR for breast and all cell markers except NAKAa1 for kidney). Nevertheless, the fact that other proteins (from the cell markers) that we probed still exhibited no statistically significant difference between healthy- and cancer tissues afford the conclusion that the normalization and protein determination steps that we have taken (all samples were delipidated during the sample-to-lysate conversions) were sufficient and appropriate. Being assured by the previous considerations, we report that cancer hotspot proteins expression were found to only weakly correlate with gene chip data from the Gene Expression Omnibus repository obtained from matched tumor-normal human breast and kidney samples. Foremost, this calls for exercising caution when one performs gene expression comparisons between healthy- and tumor tissues and extrapolates to protein expression. This should be even more concerning when one addresses tumor suppressor proteins, the ‘guardians’ of the genome. Indeed, in ^16^ it was reported that while p53 was the most differentially expressed protein between TP53 wild-type and mutated patient samples across cancer types, it intrigued the authors that in all the 11 cancer types p53 protein expression was up-regulated in the mutated group. Likewise, in ovarian tumor samples with missense hotspot mutations TP53 showed higher TP53 RNA and protein expression than those with non-hotspot mutations and without TP53 mutations ^9, 28^. Mindful of the rapid adoption of prospective clinical tumor sequencing ^29, 30^ which has led to the identification of an increasing number of somatic mutations of unknown significance, we recommend efforts quantitating protein expression of the respective gene(s) harboring those mutations to be also included. To the best of our knowledge, there is only one published attempt ^31^ to harmonize transcriptomic with proteomic analysis, but in cancer cell lines. Eventually, the harmonization of transcriptomic data with proteins expression data will assist in identifying mutations that are accompanied with significant up- or downregulation of protein expression and thus provide a better understanding of the role(s) of hotspots in cancer development.

Below we discuss our RPPA findings in light of the existing literature on the respective organ and it cancer for those oncoproteins and tumor suppressors that were statistically significantly differentially expressed in healthy-vs tumor organ tissue and there was a concomitant negative correlation between protein- and gene expression. For breast, these were: GNA11, IDH1, KIT, PDGFRA, VEGFR2, SHP2, and SMAD4, and for kidney: GNAQ, NOTCH1, SMARCB1 and TP53. For a short description regarding the function of these proteins see table 2.

### GNA11 in breast tissues

We observed a much higher GNA11 protein expression in breast tumor samples compared to matched healthy tissues, while GEO data report a decrease in gene expression. The correlation between RPPA- and Geneexp data was -0.94 (comparing means) and -0.92 (comparing medians), thus, very stark. In ^32^ it was reported that mRNA expression of GNA11 was decreased in 10 out of 16 breast cancers, and the immunoreactivity of the protein was reduced in 14 out of the 16 cancers, determined by immunohistochemistry. However, there are two potential pitfalls: i) the number of the samples examined (16) is small, mindful the large spread of both RPPA and Geneexp data among hundreds of samples (see figure 1, top right); ii) immunohistochemistry of formalin-fixed, paraffin-embedded (FFPE) samples carries a very high risk of false-positivity/negativity, plus it suffers from lack of quantitation. The differential gene expression of GNA11 in triple negative breast cancer has also been addressed in ^33^, but protein expression was not addressed.

### IDH1 in breast tissues

We observed a much higher IDH1 protein expression in breast tumor samples compared to matched healthy tissues, while GEO data report a decrease in gene expression. The correlation between RPPA- and Geneexp data was -0.83 (comparing means) and -0.80 (comparing medians), thus, also very stark. IDH2 mutational status and 2-hydroxyglutarate (2-HG) levels in serum and urine have been examined in a total of 454 female patients diagnosed with breast cancer, and results reported in ^34^. Elevation in both serum and urine 2-HG was found in patients carrying specific mutations in IDH1; however, there were no data on IDH1 protein expression or enzymatic activity or other metabolites quantified. IDH1 mutations in breast carcinomas have also been reported in ^35^, with no other data regarding the status of IDH1 gene and/or protein. IDH1-specific positive immunoreactivity in 120 out of 226 breast carcinomas examined was inversely associated with positional grading of the primary tumor (T factor), but was not compared to adjacent healthy tissue ^36^. Similarly, in ^37^ expression of IDH1 protein was only examined in breast cancer types but not healthy tissues. Finally, IDH1 gene mutations of breast tumors were addressed in ^38^, but the effect(s) at the level of protein was not examined.

### KIT in breast tissues

We observed an increase in KIT protein expression in breast tumor samples compared to matched healthy tissues, while GEO data report a decrease in gene expression. The correlation between RPPA- and Geneexp data was -0.43 (comparing means) and -0.55 (comparing medians). In the literature, the role of KIT in breast cancer is controversial: there are those supporting the concept that its loss is associated with cancer, and the other favoring that KIT is implicated in the development of breast tumors ^39^. KIT protein expression was addressed in paraffin blocks of breast tumors but no healthy samples in ^40^.

### PDGFRA in breast tissues

We observed an increase in PDGFRA protein expression in breast tumor samples compared to matched healthy tissues, while GEO data report a decrease in gene expression. The correlation between RPPA- and Geneexp data was -0.54 (comparing means) and -0.29 (comparing medians). An immunohistochemical survey of 181 formalin-fixed paraffin-embedded invasive ductal breast carcinomas revealed that PDGFRA was ‘overexpressed’, but the controls were sections of vessels in the lamina propria of gastrointestinal mucosa and blood vessels present in the periphery of the carcinomas ^41^.

### VEGFR2 in breast tissues

We observed an increase in VEGFR2 protein expression in breast tumor samples compared to matched healthy tissues (using two different antibodies), while GEO data report a decrease in gene expression. The correlation between RPPA- and Geneexp data was -0.10 – -0.34 (comparing means) and -0.17 – -0.31 (comparing medians). The expression levels of VEGFR2 has been evaluated by immunohistochemistry in PPFE blocks from breast tumors, but no comparisons to healthy tissues have been performed ^42^.

### SHP2 in breast tissues

We observed a much higher SHP2 protein expression in breast tumor samples compared to matched healthy tissues, while GEO data report a decrease in gene expression. The correlation between RPPA- and Geneexp data was -0.4 (comparing means) and -0.21 (comparing medians). GEO mining has revealed that high gene expression is associated with poorer outcome and this was in accord with protein expression assessment by immunohistochemistry: a higher SHP2 protein expression was also associated with poorer outcome ^43^, but no comparisons to healthy tissues were performed.

### GNAQ in kidney tissues

We observed a lower GNAQ protein expression in kidney tumor samples compared to matched healthy tissues, while GEO data report an increase in gene expression. The correlation between RPPA- and Geneexp data was -0.44 (comparing means) and -0.6 (comparing medians). In ^44^ it was reported that GNAQ protein expression was slightly overexpressed in tumor samples of patients with different types of renal tumors compared to matched normal samples. To the best of our knowledge, we could not find any other literature regarding GNAQ protein expression in renal cancers.

### NOTCH1 in kidney tissues

We observed a lower NOTCH1 protein expression in kidney tumor samples compared to matched healthy tissues, while GEO data report an increase in gene expression. The correlation between RPPA- and Geneexp data was -0.58 (comparing means) and -0.72 (comparing medians). NOTCH1 expression has also been evaluated by immunohistochemistry of PPFE blocks of kidney tumors in ^45^, reporting an increase in clear-cell renal carcinoma regions, compared to pericarcinoma regions and adjacent healthy tissues.

### SMARCB1 in kidney tissues

We observed a lower SMARCB1 protein expression in kidney tumor samples compared to matched healthy tissues, while GEO data report an increase in gene expression. The correlation between RPPA- and Geneexp data was -0.65 - -0.68 (comparing means) and -0.51 - -0.64 (comparing medians). SMARCB1 protein expression has been priorly investigated in FFPE samples from kidney tumors in ^46^, where endothelial/stromal cells and lymphocytes were used as controls.

### TP53 in kidney tissues

We observed a lower TP53 protein expression in kidney tumor samples compared to matched healthy tissues, while GEO data report an increase in gene expression. The correlation between RPPA- and Geneexp data was -0.52 (comparing means) and -0.56 (comparing medians). However, in the literature TP53 expression in kidney tumors is thought to be high, promoting cancer growth and metastasis, as well as a poorer outcome ^47, 48^. However, in all these studies, TP53 protein expression was addressed by immunohistochemistry of PPFE blocks and comparisons to healthy matched tissue was not performed. Apart from the counterintuitive claim that ‘overexpression’ of a tumor suppressor protein is associated with poorer outcome, the lack of proper controls (healthy matched tissues) severely confounds any conclusion.

In aggregate, the literature examined above show that in almost all cases, protein expression was performed by a means that cannot be reliably quantified (IHC of PPFE blocks) and/or healthy matched tissue controls were not available. The RPPA workflow outlined in the present work addresses the above deficiencies, in addition to demonstrating that reliance on gene expression and extrapolating to protein expression carries a very significant risk of false conclusions.

### Limitations of the study

An important limitation of the study is that the samples that we used for RPPA have not been probed for mutations of the respective genes. However, mindful that a relatively large number of gene chip data obtained from matched healthy- and tumor were fetched from NCBI-GEO (and an even large number is available for non-paired samples from the TNMplot portal), our correlations exhibit a sufficient degree of confidence. An additional limitation is that we have no patient history, nor tumor grading or subtyping of any kind, for which we could further correlate the RPPA data. Finally, mindful that RPPA is an antibody-based assay and we applied that to proteins that possibly harbor a number of mutations, it is a risk that some of these mutations would exert unknown effects on immunogenicity (and thus RPPA output signals). To this end, we have included in our antibody selection criteria the use of epitopes away from high mutagenic areas, although this was not always possible.

## Acknowledgements

We thank Prof. Várnai Péter for granting access to the Azure 600 Imaging System, Dr. Joachim Goedhart for R scripts and Dr. Leanne de Koning for providing the list of RPPA-suitable antibodies and their sources of origin. This work was supported by grants from NKFIH ([TKP2021-EGA-25], FIKP-61822-64888-EATV, VEKOP 2.3.3-15-2016-00012, 2017-2.3.4-TET-RU-2017-00003, KH129567, and K135027) to C.C.

