## Supplementary material for "RPPA survey of cancer hotspot panel proteins and cell markers in matched tumor-normal human breast and kidney samples reveals a weak correlation between proteins expression and public transcriptome repositories": Legends to supplementary figures

Legend to supplementary figure 1: A: Violin plots of the age of patients (in years) from whom the matched tumor-normal human breast and kidney samples were obtained. B: Example of dilution ratio vs protein stain data and fitted calibration curve and R^2^ that was used for sample normalization. C: Histogram of all RPPA reads of Vimentin antibody stain, indicating run (grey - Run 1, red – Run 2) D: Histogram of all RPPA reads of Vimentin antibody stain after batch normalization.

Legend to supplementary figure 2: Tumor-normal human breast sample matching (connected by a line) from each same patient fprot; blue vs red line indicates a decrease vs an increase in fprot value (healthy as control). (AB1) and (AB2) signify the type of the antibody used for the respective oncoprotein. Panel distribution matches that of main figure 1.

Legend to supplementary figure 3: Tumor-normal human breast sample matching (connected by a line) from each same patient fprot; blue vs red line indicates a decrease vs an increase in fprot value (healthy as control). (AB1) and (AB2) signify the type of the antibody used for the respective tumor suppressor. Panel distribution matches that of main figure 2.

Legend to supplementary figure 4: Tumor-normal human breast sample matching (connected by a line) from each same patient fprot; blue vs red line indicates a decrease vs an increase in fprot value (healthy as control). (AB1) and (AB2) signify the type of the antibody used for the respective cell marker. Panel distribution matches that of main figure 4.

Legend to supplementary figure 5: Tumor-normal human kidney sample matching (connected by a line) from each same patient fprot; blue vs red line indicates a decrease vs an increase in fprot value (healthy as control). (AB1) and (AB2) signify the type of the antibody used for the respective oncoprotein. Panel distribution matches that of main figure 5.

Legend to supplementary figure 6: Tumor-normal human kidney sample matching (connected by a line) from each same patient fprot; blue vs red line indicates a decrease vs an increase in fprot value (healthy as control). (AB1) and (AB2) signify the type of the antibody used for the respective tumor suppressor. Panel distribution matches that of main figure 6.

Legend to supplementary figure 7: Tumor-normal human kidney sample matching (connected by a line) from each same patient fprot; blue vs red line indicates a decrease vs an increase in fprot value (healthy as control). (AB1) and (AB2) signify the type of the antibody used for the respective cell marker. Panel distribution matches that of main figure 7.
