## Supplementary material for "RPPA survey of cancer hotspot panel proteins and cell markers in matched tumor-normal human breast and kidney samples reveals a weak correlation between proteins expression and public transcriptome repositories": Table 1

Table 1. Cancer hotspot panels offered by vendors. Source: https://app.dimensions.ai.

| ***Cancer hotspot panel name*** | ***Number of genes examined*** | ***Vendor*** | ***Number of publications**** |
| --- | --- | --- | --- |
| Ion AmpliSeq™ Cancer Hotspot Panel v2 | 50 | ThermoFisher | 6,850 |
| TruSight Oncology 500 | 523 | Illumina | 578 |
| CleanPlex OncoZoom Cancer Hotspot Kit | 65 | Paragon Genomics | 230 |
| Tapestri Single-cell DNA Tumor Hotspot Panel | 59 | Mission Bio | 334 |
| NEBNext Direct® Cancer HotSpot Panel | 50 | New England Biolabs | 1,045 |
| OncoGxOne™ Discovery cancer panels | 150-400 | GENEWIZ | 10 |
| Azenta Pan-Cancer Panel | 634 | Azenta Life Sciences | 236 |
| Cancer Hotspot Panel | 65 | CD Genomics | 524 |

* publications: datasets, grants, patents, clinical trials, policy documents (mined out of 139,456,791 entries as of October 2023)
