## Supplementary material for "RPPA survey of cancer hotspot panel proteins and cell markers in matched tumor-normal human breast and kidney samples reveals a weak correlation between proteins expression and public transcriptome repositories": Table 2

Table 2. Name of proteins represented by the Ion AmpliSeq™Cancer Hotspot Panel v2 tested by RPPA, their nomenclature, short description, Research Resource Identifiers (RRIDs) of antibodies used, epitopes (epitopes for invalid antibodies are not provided) and titers. RRIDs in **bold** signify the valid antibodies used for RPPA; if not in bold, it means that they were tested but found not to be suitable due to either lack of monospecificity or no labelling or band appeared far from the expected MW range, in Western blotting. When two antibodies are denoted in **bold** (for a single protein, thus both yielded suitable labelling) they are arbitrarily additionally annotated as **AB1** or **AB2** to distinguish them in the text and figures. No suitable antibody was found for PIK3CA and SMO proteins.

| **Name of protein** | **Aliases** | **Abbreviation** | **UniProt** | **Gene** | **Short description of protein function** | **RRID** | **Epitope** | **Titer** |
| --- | --- | --- | --- | --- | --- | --- | --- | --- |
| Tyrosine-protein kinase ABL1 | Proto-oncogene c-Abl; Abelson murine leukemia viral oncogene homolog 1; Abelson tyrosine-protein kinase 1; p150 | ABL1 | P00519 | *ABL1 (Synonym: ABL, JTK7)* | A non-receptor tyrosine-protein kinase belonging to the Abelson (ABL) family of protein kinases; plays a role in processes linked to cytoskeleton remodeling in response to extracellular stimuli, cell motility and adhesion, receptor endocytosis, autophagy, DNA damage response, apoptosis and in the phosphorylation of multiple receptor tyrosine kinases promoting endocytosis of EGFR. It has also been reported to be a tumor suppressor protein. | **RRID:AB_2257757** | Residues surrounding Pro580 | 1:1,000 |
| RAC-alpha serine/threonine-protein kinase | Protein kinase B alpha; Proto-oncogene c-Akt;  RAC-PK-alpha | AKT1 | P31749 | *AKT1*  *(PKB, RAC)* | A serine/threonine kinase previously known as protein kinase B (PKB) involved in cellular survival pathways by inhibiting apoptotic processes. | **RRID:AB_915783** | Residues near the C-terminus; vendor claims it reacts with all three isoforms, but the validator IC reported it to react only with AKT1 | 1:1,000 |
| ALK tyrosine kinase receptor | Anaplastic lymphoma kinase;  CD246 | ALK | Q9UM73 | *ALK* | A receptor tyrosine kinase activating multiple pathways through phospholipase C, JAK, STAT, PI3K/Akt, mTOR and MAPK signaling cascades. | **RRID:AB_11127207** | Residues in the C-terminus | 1:5,000 |
| Adenomatous polyposis coli protein | Protein APC | APC | P25054 | *APC*  *(DP2.5)* | Tumor suppressor protein. It performs multiple functions through WNT-dependent (promoting β-catenin phosphorylation, ubiquitination and subsequent proteolytic degradation) and WNT-independent mechanisms (contributes to apical-basal polarity, microtubule networks, cell cycle progression, DNA replication and repair, apoptosis, and cell migration). | **RRID:AB_628735 (AB1)** | Residues near the C-terminus | 1:1,000 |
|  |  |  |  |  |  | **RRID:AB_2057497 (AB2)** | Residues near the N-terminus | 1:2,000 |
| Serine-protein kinase ATM | Ataxia telangiectasia mutated | ATM | Q13315 | *ATM*  *(TEL1, TELO1)* | Belongs to the phosphatidylinositol 3-kinase-related kinase (PIKK) super family; upon activation it phosphorylates multiple proteins involved in DNA repair. It is also involved in cell cycle checkpoints, activates p53 and apoptosis and mediates mitochondrial homeostasis and reactive oxygen species regulation. Frequently considered as a tumor suppressor protein. | **RRID:AB_725574** | Proprietary | 1:1,000 |
| Serine/threonine-protein kinase B-raf | Proto-oncogene B-Raf; p94 | BRAF | P15056 | *BRAF* | Belongs to the rapidly accelerated fibrosarcoma (RAF) family of serine/threonine kinases and functions as a Mitogen-activated pathway kinase kinase kinase (MAPKKK) in the MAP kinase/ERK signaling cascade leading to direct and indirect transcriptional regulation involved in cell survival and proliferation. | **RRID:AB_725762** | Proprietary | 1:1,000 |
| Cadherin-1 | CAM 120/80; CD324;  Epithelial cadherin; E-cadherin; Uvomorulin | CDH1 | P12830 | *CDH1*  *(CDHE, UVO)* | Tumor suppressor protein. In normal epithelial tissues mature E-cadherin binds polarized epithelial cells together at lateral surface *via* adherens junctions and complexes with β-catenin preventing it from being released into the cytoplasm and entering the nucleus, thus preventing activation of the WNT signaling pathway. | **RRID:AB_731493** | Amino acids 604-622 | 1:50,000 |
| Cyclin-dependent kinase inhibitor 2A | Cyclin-dependent kinase 4 inhibitor A; CDK4I; p16-INK4a; Multiple tumor suppressor 1 | p16INK4A | P42771 | ***CDKN2A***  *(CDKN2, MTS1)* | The CDKN2A locus encodes two gene products: p16INK4A and p14ARF, each being under transcriptional regulation by independent promoters and each with distinct tumor suppressive functions, thus, p16INK4A is a tumor suppressor protein. It is a cyclin-dependent kinase (CDK) inhibitor that binds to CDK4 and CDK6 and prevents the phosphorylation of retinoblastoma protein (RB1), prohibiting cell cycle progression. | **RRID:AB_10858268 (AB1)** | Within 100 residues to the C-terminus | 1:10,000 |
|  |  |  |  |  |  | **RRID:AB_2750891 (AB2)** | Residues surrounding Ala34 | 1:1,000 |
|  |  |  |  |  |  | RRID:AB_1640753 |  |  |
| Tumor suppressor ARF | Tumor suppressor ARF; Alternative reading frame; ARF; p14ARF; Cyclin-dependent kinase inhibitor 2A | p14ARF | Q8N726 | ***CDKN2A*** | Tumor suppressor protein. p14ARF specifically binds to HDM2, a protein with the E3 ubiquitin ligase activity and promotes its degradation, thereby blocking HDM2-induced degradation of p53 and enhancing p53-dependent transactivation and apoptosis. | **RRID:AB_2934126** | Within 100 residues to the C-terminus (different from AB_10858268) | 1:1,000 |
|  |  |  |  |  |  | RRID:AB_2934127 |  |  |
| Macrophage colony-stimulating factor 1 receptor | Proto-oncogene c-Fms; M-CSF-R; CD115 | CSF1R | P07333 | *CSF1R*  *(FMS)* | A tyrosine kinase transmembrane receptor and member of the CSF1/PDGF receptor family of tyrosine-protein kinases. By binding its ligand, colony stimulating factor 1, it controls the production, differentiation, and function of hematopoietic precursor cells, especially mononuclear phagocytes, such as macrophages and monocytes. In some cancers and under some conditions, it is considered a tumor suppressor protein. | **RRID:AB_2934128** | Residues within amino acids 300-550. | 1:2,000 |
|  |  |  |  |  |  | RRID:AB_2934129 |  |  |
| Catenin beta-1 | Beta-catenin | CTNNB1 | P35222 | *CTNNB1* | In the absence of WNT, beta-catenin forms a complex which is ubiquitinated and degraded, while in the presence of WNT ligand, CTNNB1 is not ubiquitinated and accumulates in the nucleus acting as a coactivator for transcription factors of the TCF/LEF family, leading to the activation of WNT-responsive genes. | **RRID:AB_823447** | Residues near the C-terminus | 1:1,000 |
| Epidermal growth factor receptor | Proto-oncogene c-ErbB-1; Receptor tyrosine-protein kinase erbB-1 | EGFR | P00533 | *EGFR*  *(ERBB, ERBB1, HER1)* | A transmembrane glycoprotein that constitutes one of four members of the erbB family of tyrosine kinase receptors activating several signaling cascades, such as the RAS-RAF-MEK-ERK, PI3 kinase-AKT, PLCgamma-PKC and STATs leading to cell proliferation. | **RRID:AB_2246311** | Residues forming the cytoplasmic domain | 1:1,000 |
| Receptor tyrosine-protein kinase erbB-2 | Proto-oncogene Neu; Proto-oncogene c-ErbB-2; Tyrosine kinase-type cell surface receptor HER2; p185erbB2; CD340 | ERBB2 | P04626 | *ERBB2*  *(HER2, MLN19, NEU, NGL)* | Member of the epidermal growth factor (EGF) receptor family of receptor tyrosine kinases but it has no ligand binding domain of its own, therefore heterodimerisation with other ligand-bound EGF receptor family members is required to stabilize their ligand binding thereby enhancing kinase-mediated activation of downstream signaling pathways, such as those involving mitogen-activated protein kinase and phosphatidylinositol-3 kinase. Considered to be a tumor suppressor protein. | **RRID:AB_2099242** | Residues near the C-terminus | 1:1,000 |
| Receptor tyrosine-protein kinase erbB-4 | Tyrosine kinase-type cell surface receptor HER4; c-ErbB-4; HER4; p180erbB4 | ERBB4 | Q15303 | *ERBB4*  *(HER4)* | Member of the Tyr protein kinase family and the epidermal growth factor receptor subfamily and is required for normal development of the heart, the central nervous system and the mammary gland, for gene transcription, cell proliferation, differentiation, migration and apoptosis. It has been proposed to act as an oncogene product and tumor suppressor protein. | **RRID:AB_1566046** | Proprietary | 1:1,000 |
| Histone-lysine N-methyltransferase EZH2 | ENX-1; Enhancer of zeste homolog 2 | EZH2 | Q15910 | *EZH2*  *(KMT6)* | Regulator of transcription as a member of polycomb repressive complex 2 (PRC2) which exerts repression of downstream genes over successive cell generations; its abnormal expression is correlated to invasiveness and progression of different tumors, for example by induction of epithelial-mesenchymal transition (EMT). It has been proposed to act as an oncogene product and tumor suppressor protein. | **RRID:AB_10694683** | Residues surrounding Arg354 | 1:5,000 |
|  |  |  |  |  |  | RRID:AB_2934130 |  |  |
| F-box/WD repeat-containing protein 7 | F-box and WD-40 domain-containing protein 7; F-box protein FBX30; SEL-10 | FBXW7 | Q969H0 | *FBXW7*  *(FBW7, FBX30, SEL10)* | A tumor suppressor protein that recognizes substrates for ubiquitin-mediated degradation through the presence of a conserved CDC4 phosphodegron (CPD) motif. | **RRID:AB_1840617** | 109 residues to the C-terminus | 1:500 |
|  |  |  |  |  |  | RRID:AB_2626551 |  |  |
|  |  |  |  |  |  | RRID:AB_1078484 |  |  |
|  |  |  |  |  |  | RRID:AB_2724297 |  |  |
| Fibroblast growth factor receptor 1 | Basic fibroblast growth factor receptor 1; BFGFR; bFGF-R-1; Fms-like tyrosine kinase 2; FLT-2; N-sam; Proto-oncogene c-Fgr; CD331 | FGFR1 | P11362 | *FGFR1*  *(BFGFR, CEK, FGFBR, FLG, FLT2, HBGFR)* | A subset of the Receptor Tyrosine Kinase (RTK) family is the  fibroblast growth factor receptor (FGFR) family, which contains  four homologous receptors: FGFR1, FGFR2, FGFR3, and FGFR4. Alternative splicing of 8–9 exons generate two different isoforms in each of FGFR1, FGFR2 and FGFR3 except for the FGFR4 gene, which does not show any splicing activity, thus there are 7 different FGF receptors. The members differ from one another in their ligand affinities and tissue distribution. FGFR signaling results in cellular proliferation and migration, anti-apoptosis, angiogenesis and wound healing. | **RRID:AB_11178519** | Residues near the C-terminus | 1:4,000 |
|  |  |  |  |  |  | RRID:AB_1523613 |  |  |
| Fibroblast growth factor receptor 2 | K-sam; KGFR; Keratinocyte growth factor receptor; CD332 | FGFR2 | P21802 | *FGFR2*  *(BEK, KGFR, KSAM)* | Mutations leading to constitutive kinase activation or impair normal FGFR2 maturation, internalization and degradation lead to aberrant signaling in several tumor types, while amplification occur frequently in breast cancer, predominantly the triple-negative type. It has been proposed to act as an oncogene product and tumor suppressor protein. | **RRID:AB_2797742** | Residues surrounding amino acid 440. The protein exhibits 17 isoforms; several amino acids can occupy position 440; it is not known which was used for generating AB_2797742. | 1:2,000 |
|  |  |  |  |  |  | RRID:AB_2934131 |  |  |
| Fibroblast growth factor receptor 3 | CD333 | FGFR3 | P22607 | *FGFR3*  *(JTK4)* | Plays a role in bone development and maintenance; mutations that lead to constitutive kinase activation or impair normal FGFR3 maturation, internalization and degradation lead to aberrant signaling. Depending on cancer type, . It was shown to act as either an oncogene product or tumor suppressor protein. | **RRID:AB_2246903** | Residues forming the cytoplasmic domain | 1:3,000 |
|  |  |  |  |  |  | RRID:AB_627596 |  |  |
| Receptor-type tyrosine-protein kinase FLT3 | FL cytokine receptor; Fetal liver kinase-2; FLK-2;  Fms-like tyrosine kinase 3; FLT-3; Stem cell tyrosine kinase 1; STK-1;  CD135 | FLT3 | P36888 | *FLT3*  *(CD135, FLK2, STK1)* | A tyrosine kinase promoting the activation of downstream pathways involving phosphatidylinositol-3 kinase (PI3K), AKT, mammalian target of rapamycin (mTOR), RAS, and extracellular signal-related kinase (ERK) thereby regulating differentiation, proliferation and survival of hematopoietic progenitor cells and dendritic cells. | **RRID:AB_2049358** | Proprietary | 1:1,000 |
|  |  |  |  |  |  | RRID:AB_2107052 |  |  |
| Guanine nucleotide-binding protein subunit alpha-11 | Guanine nucleotide-binding protein G(y) subunit alpha | GNA11 | P29992 | *GNA11*  *(GA11)* | Subunit alpha-11 of guanine nucleotide-binding proteins (G proteins) signaling to phospholipase C. | **RRID:AB_2715558** | Amino acids 3-39 | 1:500 |
| Guanine nucleotide-binding protein G(s) subunit alpha isoforms short | Adenylate cyclase-stimulating G alpha protein | GNAS | P63092 | *GNAS* | GNAS (Guanine Nucleotide binding protein, Alpha Stimulating) is a protein product of a complex imprinted gene that generates multiple gene products through the use of multiple promoters and first exons that splice onto a common set of downstream exons (exons 2–13). The major GNAS gene product, which is generated by the most downstream promoter (exon 1), is the ubiquitously expressed G protein α-subunit (Gsα, Uniprot: P63092) that links numerous hormonal and other seven-transmembrane receptors to adenylyl cyclase and is required for the receptor-stimulated intracellular cAMP production. The most upstream GNAS promoter generates transcripts for the chromogranin-like protein NESP55 (Uniprot: O95467), which is structurally and functionally unrelated to Gsα. The next promoter generates transcripts encoding the neuroendocrine-specific Gsα isoform XLαs (Uniprot: Q5JWF2), which is structurally identical to Gsα, except for the presence of an extra-long amino-terminal extension encoded by its specific first exon. XLas isoforms interact with the same set of receptors as GNAS isoforms. In some cancers, GNAS is a tumor suppressor protein. | **RRID:AB_2934132** | Amino acids 42-188 | 1:1,000 |
|  |  |  |  |  |  | RRID:AB_2934133 |  |  |
| Guanine nucleotide-binding protein G(q) subunit alpha | Guanine nucleotide-binding protein alpha-q | GNAQ | P50148 | *GNAQ*  *(GAQ)* | Alpha subunit of a heterotrimeric Guanine nucleotide binding protein that mediates stimulation of phospholipase C-beta. | **RRID:AB_2934134** | Residues near the C-terminus | 1:4,000 |
| Hepatocyte nuclear factor 1-alpha | Liver-specific transcription factor LF-B1; Transcription factor 1; TCF-1 | HNF1A | P20823 | *HNF1A*  *(TCF1)* | Belongs to the homeobox protein family and is an essential transcription factor for many hepatic genes - involved in detoxification, homeostasis and metabolisms of glucose, lipid, steroid and amino acid - and embryonic pancreatic development and differentiation as well as maintaining the growth and function of islet β cells. A potential tumor suppressor protein. | **RRID:AB_2538735** | Amino acids 35-268 | 1:1,000 |
|  |  |  |  |  |  | RRID:AB_2728751 |  |  |
|  |  |  |  |  |  | RRID:AB_2279592 |  |  |
|  |  |  |  |  |  | RRID:AB_2934143 |  |  |
| GTPase HRas | c-H-ras; Transforming protein p21; p21ras | HRAS | P01112 | *HRAS*  *(HRAS1)* | Ras, an important family of proto-oncogenes in humans, consists of three members (H-Ras (Harvey-RAS), N-Ras and K-Ras) and they are involved in signal transduction pathways (MAPK, PI3K) due to their intrinsic GTPase activity which is related to a continuous cycle of de- and re-palmitoylation, in turn regulating its rapid exchange between the plasma membrane and the Golgi apparatus. Only the endoplasmic reticulum-associated form of HRAS can activate the RAF1-ERK signaling pathway, leading to fibroblast transformation; the Golgi-associated form seems to be unable to induce cell transformation or proliferation.  Mutations in HRAS can be found in dermatological malignancies and head and neck cancers | **RRID:AB_1090245** | Amino acids 92-108 | 1:4,000 |
| Isocitrate dehydrogenase [NADP] cytoplasmic | Cytosolic NADP-isocitrate dehydrogenase;  IDPc; NADP(+)-specific ICDH; Oxalosuccinate decarboxylase | IDH1 | O75874 | *IDH1*  *(PICD)* | NADP^+^-dependent isocitrate dehydrogenase found in the cytoplasm and peroxisomes and catalyzes the NADP^+^-dependent oxidative decarboxylation of isocitrate (D-threo-isocitrate) to alpha-ketoglutarate (2-oxoglutarate). | **RRID:AB_2864315** | Residues within 1-100 from the N-terminus | 1:1,000 |
| Isocitrate dehydrogenase [NADP], mitochondrial | NADP(+)-specific ICDH; Oxalosuccinate decarboxylase;  ICD-M | IDH2 | P48735 | *IDH2* | Mitochondrial isoform, catalyzes the NAD^+^-dependent oxidative decarboxylation of isocitrate (D-threo-isocitrate) to alpha-ketoglutarate (2-oxoglutarate). | **RRID:AB_2799511** | Residues surrounding Val195 | 1:4,000 |
| Tyrosine-protein kinase JAK2 | Janus kinase 2 | JAK2 | O60674 | *JAK2* | A non-receptor tyrosine kinase expressed widespread in human cells mediating essential signaling events in both innate and adaptive immunity and growth factors leading to regulation of proliferation, differentiation, migration, apoptosis and cell survival. May also act as a tumor suppressor protein. | **RRID:AB_2128522** | Residues surrounding Pro841 | 1:1,000 |
| Tyrosine-protein kinase JAK3 | Janus kinase 3; Leukocyte janus kinase; L-JAK | JAK3 | P52333 | *JAK3* | A non-receptor tyrosine kinase specifically associated with cytokine receptors and it is expressed in both immune cells and in intestinal epithelial cells (IECs), therefore activating Jak3 mutations lead to the development of hematologic and epithelial cancers. | **RRID:AB_10999548** | Residues near the C-terminus | 1:4,000 |
|  |  |  |  |  |  | RRID:AB_775811 |  |  |
| Vascular endothelial growth factor receptor 2 | VEGFR-2; Fetal liver kinase 1; FLK-1; Kinase insert domain receptor; KDR; CD309 | VEGFR2 | P35968 | *KDR*  *(FLK1, VEGFR2)* | A tyrosine-protein kinase that acts as a cell-surface receptor for VEGF(A, C and D) playing a role in regulation of angiogenesis, vascular development, vascular permeability and embryonic hematopoiesis; it promotes proliferation, survival, migration and differentiation of endothelial cells. | **RRID:AB_11178792 (AB1)** | 150 residues to the C-terminus | 1:2,000 |
|  |  |  |  |  |  | **RRID:AB_2212507 (AB2)** | 150 residues to the C-terminus; different clone from AB_11178792 | 1:2,000 |
|  |  |  |  |  |  | RRID:AB_2934135 |  |  |
| Mast/stem cell growth factor receptor Kit | SCFR; Proto-oncogene c-Kit; v-kit Hardy-Zuckerman 4 feline sarcoma viral oncogene homolog; CD117 | KIT | P10721 | *KIT*  *(SCFR)* | A receptor tyrosine kinase that once activated by its cytokine ligand, stem cell factor (SCF), plays a role in the proliferation, differentiation, migration and apoptosis of many cell types involved in hematopoiesis; also involved in stem cell maintenance, mast cell development, melanogenesis and gametogenesis. | **RRID:AB_2799120** | Residues surrounding Val955 | 1:4,000 |
|  |  |  |  |  |  | RRID:AB_731513 |  |  |
| GTPase KRas | c-K-ras; Ki-Ras | KRAS | P01116 | *KRAS*  *(KRAS2, RASK2)* | Encoded by a gene that belongs to the Kirsten ras oncogene homolog from the mammalian ras gene family, is a member of the small GTPase superfamily (HRAS, KRAS and NRAS) and it is the most oncogenic with its 85% share of all mutated RAS proteins observed in cancer. | **RRID:AB_2797685** | Proprietary | 1:4,000 |
|  |  |  |  |  |  | RRID:AB_2532192 |  |  |
| Hepatocyte growth factor receptor (HGF) | Proto-oncogene c-Met; Scatter factor receptor | MET | P08581 | *MET* | A type of receptor tyrosine kinase that is expressed on the surfaces of various epithelial cells; its ligand is HGF/SF(ligand hepatocyte growth factor/scatter factor) and under normal conditions, c-Met mediates embryogenesis, tissue regeneration, wound healing, and the formation of nerve and muscle tissue which is controlled by the tumor suppressor p53; abnormal activation of c-Met can promote the development and progression of multiple cancers. | **RRID:AB_10858224** | Residues near the C-terminus | 1:1,000 |
| DNA mismatch repair protein Mlh1 | MutL protein homolog 1 | MLH1 | P40692 | *MLH1*  *(COCA2)* | This protein can heterodimerize with mismatch repair endonuclease PMS2 to form a complex with an endogenous endonuclease activity that incises the unmethylated strand; the single-strand breaks generated by this manner signals for downstream repair processes; it can also heterodimerize with DNA mismatch repair protein MLH3 to form MutL gamma, which is involved in meiosis. A tumor suppressor protein. | **RRID:AB_2049968** | Amino acids 700-800 | 1:10,000 |
|  |  |  |  |  |  | RRID:AB_2145615 |  |  |
|  |  |  |  |  |  | RRID:AB_10545447 |  |  |
| Thrombopoietin receptor | Myeloproliferative leukemia protein; Proto-oncogene c-Mpl; CD110 | TPOR | P40238 | *MPL*  *(TPOR)* | Expressed predominantly on the surface of megakaryocytes, platelets, hemangioblasts and hematopoietic stem cells, TPOR is the receptor of the cytokine thrombopoietin (TPO) and is a chief regulator of megakaryocyte and platelet production through the Janus kinase (JAK) family. | **RRID:AB_10858953** | 600 residues to the C-terminus | 1:4,000 |
| Neurogenic locus notch homolog protein 1 | Translocation-associated notch protein TAN-1 | NOTCH1 | P46531 | *NOTCH1*  *(TAN1)* | The preproprotein is proteolytically processed in the trans-Golgi network to generate two polypeptide chains that heterodimerize to form the mature cell-surface receptor and its regulated signaling is critical for the generation of normal thymic T-cells, remaining crucial after the release of T-cells into the periphery, therefore abnormal Notch signaling results in leukemia. Considered both an oncogene and a tumor suppressor, depending on conditions. | **RRID:AB_2153354** | Residues surrounding Pro2438 | 1:1,000 |
|  |  |  |  |  |  | RRID:AB_881725 |  |  |
| Nucleophosmin | Nucleolar phosphoprotein B23; Nucleolar protein NO38; Numatrin | NPM1 | P06748 | *NPM1* | Belongs to a histone chaperones family; is an abundant nucleolar protein found in proliferating cells. Upon oligomerization it exerts an effect on cellular proliferation, while the monomeric form is associated with having a role in the DNA damage response and induction of apoptosis; Considered both an oncogene and a tumor suppressor, depending on conditions. | **RRID:AB_2934136 (AB1)** | 200 residues to the C-terminus | 1:1,000 |
|  |  |  |  |  |  | **RRID:AB_881735 (AB2)** | Amino acids 1-100 | 1:200,000 |
| GTPase NRas | Transforming protein N-Ras | NRAS | P01111 | *NRAS* | Neuroblastoma rat sarcoma (RAS) viral oncogene homolog (NRAS), a small GTPase; it controls cell growth, differentiation, and survival by facilitating signal transduction by shuttling between the Golgi apparatus and the plasma membrane regulated through palmitoylation and depalmitoylation. Constitutively active NRAS mutants signal through several pathways – most notably, the RAS–RAF–MAPK and PI3K–AKT pathways and induce cell-cycle dysregulation, pro-survival pathways, and cellular proliferation. | **RRID:AB_2934137** | Amino acids 70-101 | 1:1,000 |
| Platelet-derived growth factor receptor alpha | Alpha-type platelet-derived growth factor receptor; CD140a antigen;  Platelet-derived growth factor receptor 2; PDGFR-2 | PDGFRA | P16234 | *PDGFRA*  *(PDGFR2, RHEPDGFRA)* | A tyrosine-protein kinase that acts as a cell-surface receptor for PDGFA, -B and -C and plays an essential role in the regulation of embryonic development and depending on the context, promotes or inhibits cell proliferation, survival and cell migration. | **RRID:AB_2162345** | Residues near the C-terminus | 1:1,000 |
| Phosphatidylinositol 4,5-bisphosphate 3-kinase catalytic subunit alpha isoform | Phosphoinositide 3-kinase alpha;  Phosphoinositide-3-kinase catalytic alpha polypeptide; Serine/threonine protein kinase PIK3CA | PIK3CA | P42336 | *PIK3CA* | PIK3CA is the catalytic subunit of phosphatidylinositol 3-kinase which phosphorylates phosphatidylinositol (PI); its phosphorylated derivatives activate signaling cascades involved in cell growth, survival, proliferation, motility and morphology in response to various growth factors, such as EGF, insulin, IGF1, VEGFA and PDGF. | RRID:AB_2165248 | *No suitable Ab found* |  |
|  |  |  |  |  |  | RRID:AB_2934138 |  |  |
|  |  |  |  |  |  | RRID:AB_777253 |  |  |
| Phosphatidylinositol 3,4,5-trisphosphate 3-phosphatase and  dual-specificity protein phosphatase PTEN | Phosphatase and tensin homolog;  Mutated in multiple advanced cancers 1 | PTEN | P60484 | *PTEN*  *(MMAC1, TEP1)* | Tumor suppressor protein. It is a phosphatidylinositol-3,4,5-trisphosphate 3-phosphatase negatively regulating AKT/PKB signaling pathway; also acting as a dual-specificity protein phosphatase dephosphorylating tyrosine-, serine- and threonine-phosphorylated proteins. | **RRID:AB_390810** | Residues near the C-terminus | 1:1,000 |
| Tyrosine-protein phosphatase non-receptor type 11 | SH-PTP2; SH-PTP3;  Protein-tyrosine phosphatase 1D; PTP-1D; Protein-tyrosine phosphatase 2C; PTP-2C | SHP2 | Q06124 | *PTPN11 (SHP2,*  *PTP2C, SHPTP2)* | Src homology 2 domain-containing protein tyrosine phosphatase 2 (Shp2) is a ubiquitously expressed protein tyrosine phosphatase (PTP) which is normally self-inhibited; upon activation (by the presence of phosphorylated tyrosine (pY) residues) it exerts a positive role in transducing signals initiated from receptor and cytosolic kinases; in addition to growth factor/cytokine-induced signaling, Shp2 regulates genotoxic stress triggered signaling and cellular responses. Predominantly oncogenic, but can be tumor suppressor in a context-dependent manner. | **RRID:AB_779072** | Amino acids 1-100 | 1:4,000 |
| Retinoblastoma-associated protein | p105-Rb; p110-RB1;  pRb; Rb; pp110 | RB1 | P06400 | *RB1* | Tumor suppressor protein. Regulator of the G1/S transition of the cell cycle also promoting the G0-G1 transition upon phosphorylation and activation by CDK3/cyclin-C. | **RRID:AB_823629** | Residues surrounding His890 | 1:2,000 |
| Proto-oncogene tyrosine-protein kinase receptor Ret | Cadherin family member 12; Proto-oncogene c-Ret | RET | P07949 | *RET*  *(CDHF12, CDHR16, PTC)* | A receptor tyrosine-protein kinase proto-oncogene involved in cell proliferation, neuronal navigation, cell migration, and cell differentiation which can undergo oncogenic activation through both cytogenetic rearrangement and point mutations. Oncogene protein and a potential tumor suppressor protein, depending on conditions. | **RRID:AB_2238465 (AB1)** | Proprietary | 1:1,000 |
|  |  |  |  |  |  | **RRID:AB_2798509 (AB2)** | Residues surrounding Pro320 | 1:2,000 |
|  |  |  |  |  |  | RRID:AB_2049540 |  |  |
|  |  |  |  |  |  | RRID:AB_2920824 |  |  |
| Mothers against decapentaplegic homolog 4 | Deletion target in pancreatic carcinoma 4; SMAD family member 4 | SMAD4 | Q13485 | *SMAD4*  *(DPC4, MADH4)* | Tumor suppressor protein. SMAD4 is a member of the Smad family of signal transduction proteins, which is activated by transmembrane serine-threonine receptor kinases in response to transforming growth factor (TGF)-beta signaling and accumulate in the nucleus where it regulates the transcription of target genes as a component of multimeric complexes inhibiting epithelial cell proliferation. | **RRID:AB_2728776** | Residues surrounding Asp165 | 1:1,000 |
| SWI/SNF-related matrix-associated actin-dependent regulator of chromatin  subfamily B member 1 | BRG1-associated factor 47; Integrase interactor 1 protein; SNF5 homolog | SMARCB1 | Q12824 | *SMARCB1*  *(BAF47, INI1, SNF5L1)* | Tumor suppressor protein. Core component of the BAF (hSWI/SNF) complex that relieves repressive chromatin structures, allowing the transcriptional machinery to access its targets, therefore playing important roles in cell proliferation and differentiation and cellular antiviral mechanisms. | **RRID:AB_2715495 (AB1)** | Residues near the C-terminus | 1:50,000 |
|  |  |  |  |  |  | **RRID:AB_2934139 (AB2)** | Residues near the C-terminus (different from AB_2715495) | 1:10,000 |
| Smoothened homolog | Protein Gx | SMO | Q99835 | *SMO*  *(SMOH)* | SMO is a G protein-coupled receptor that potentially associates with the patched protein (PTCH) in order to transduce the hedgehog's proteins signal. | RRID:AB_2239686 | *No suitable Ab found* |  |
|  |  |  |  |  |  | RRID:AB_2934144 |  |  |
|  |  |  |  |  |  | RRID:AB_2934145 |  |  |
| Proto-oncogene tyrosine-protein kinase Src | Proto-oncogene c-Src; pp60c-src | SRC | P12931 | *SRC*  *(SRC1)* | A non-receptor protein tyrosine kinase proto-oncogene, highly similar to the v-src gene of Rous sarcoma virus, participating in signaling pathways that control gene transcription, immune responses, cell adhesion, cell cycle progression, apoptosis, migration, and transformation. | **RRID:AB_2106047 (AB1)** | Amino acids 1-100 | 1:2,000 |
|  |  |  |  |  |  | **RRID:AB_10865528 (AB2)** | Amino acids 1-100 (different from AB_2106047) | 1:50,000 |
| Serine/threonine-protein kinase STK11 | Liver kinase B1; LKB1; Renal carcinoma antigen NY-REN-19 | STK11 | Q15831 | *STK11 (LKB1)* | Tumor suppressor protein. A serine/threonine kinase that regulates cell polarity (by remodeling the actin cytoskeleton) and energy metabolism (through AMPK family members), apoptosis (mediator of p53/TP53-dependent apoptosis) and DNA damage response. | **RRID:AB_2889913** | Residues within amino acids 1-350 | 1:1,000 |
| Cellular tumor antigen p53 | Antigen NY-CO-13;  Phosphoprotein p53;  Tumor suppressor p53 | TP53 | P04637 | *TP53* | Tumor suppressor protein. | **RRID:AB_10695803** | Proprietary | 1:1,000 |
| von Hippel-Lindau disease tumor suppressor | Protein G7; pVHL | VHL | P40337 | *VHL* | Tumor suppressor protein. A component of a ubiquitination complex involved in the degradation of hypoxia-inducible-factor (HIF); also playing a role in cytokine signaling, regulation of senescence, and formation of the extracellular matrix; variants of this gene are associated with von Hippel-Lindau syndrome, pheochromocytoma, renal cell carcinoma, and cerebellar hemangioblastoma. | **RRID:AB_2879705** | Amino acids 90-172 | 1:500 |
|  |  |  |  |  |  | RRID:AB_10807698 |  |  |
|  |  |  |  |  |  | RRID:AB_2934140 |  |  |
