## Supplementary material for "RPPA survey of cancer hotspot panel proteins and cell markers in matched tumor-normal human breast and kidney samples reveals a weak correlation between proteins expression and public transcriptome repositories": Table 3

Table 3. Name of cell markers used in RPPA, abbreviations (used in the text and figures), Research Resource Identifiers (RRIDs) of antibodies used and their titers.

| **Name of protein** | **Abbreviation** | **RRID** | **Titer** |
| --- | --- | --- | --- |
| α-tubulin | TUBA1A | AB_2619646 | 1:4,000 |
| β-actin | ACTB | AB_2223210 | 1:10,000 |
| Calreticulin | CALR | AB_2688013 | 1:4,000 |
| Caveolin-1 | CAV1 | AB_2275453 | 1:4,000 |
| Desmin | DES | AB_1903947 | 1:4,000 |
| Fibrillarin | FBL | AB_2278087 | 1:4,000 |
| Golgi matrix protein 130 | GOLGA2 | AB_2797933 | 1:4,000 |
| Insulin-like growth factor II | IGF2R | AB_2798462 | 1:4,000 |
| Keratin 19 | KRT19 | AB_2722626 | 1:4,000 |
| Lysine-specific histone demethylase 1A | LSD1 | AB_2070132 | 1:4,000 |
| Na^+^/K^+^ ATPase α1 subunit | NAKAa1 | AB_2798866 | 1:4,000 |
| Nuclear pore complex protein Nup98-Nup96 | NUP98 | AB_2267700 | 1:4,000 |
| Ribosomal Protein S3 | RPS3 | AB_10622028 | 1:4,000 |
| Vimentin | VIM | AB_10695459 | 1:4,000 |
